# Parallel evolution and enhanced virulence upon *in vivo* passage of an RNA virus in *Drosophila melanogaster*

**DOI:** 10.1101/2023.07.21.549997

**Authors:** Oscar M. Lezcano, Lara Fuhrmann, Gayatri Ramakrishnan, Niko Beerenwinkel, Martijn A. Huynen, Ronald P. van Rij

**Affiliations:** Department of Medical Microbiology, Radboud University Medical Center, P.O. Box 9101, 6500 HB, Nijmegen, the Netherlands; Department of Biosystems Science and Engineering, ETH Zurich, Basel, 4058, Switzerland; SIB Swiss Institute of Bioinformatics, Basel, 4058, Switzerland; Department of Medical BioSciences, Radboud University Medical Center, P.O. Box 9101, 6500 HB, Nijmegen, the Netherlands

**Keywords:** Drosophila C virus, RNA interference, Dicer, insect immunity, virus evolution

## Abstract

Virus evolution is strongly affected by antagonistic co-evolution of virus and host. Host immunity positively selects for viruses that evade the immune response, which in turn may drive counter-adaptations in host immune genes. We investigated how host immune pressure shapes virus populations, using the fruit fly *Drosophila melanogaster* and its natural pathogen Drosophila C virus (DCV), as a model. We performed an experimental evolution study in which DCV was serially passaged for ten generations in three fly genotypes differing in their antiviral RNAi response: wild-type flies and flies in which the endonuclease gene *Dicer-2* was either overexpressed or inactivated. All evolved virus populations replicated more efficiently *in vivo* and were more virulent than the parental stock. The number of polymorphisms increased in all three host genotypes with passage number, which was most pronounced in *Dicer-2* knockout flies. Mutational analysis showed strong parallel evolution, as mutations accumulated in a specific region of the VP3 capsid protein in every lineage in a host genotype-independent manner. The parental tyrosine at position 95 of VP3 was substituted with either one of five different amino acids in 14 out of 15 lineages. However, no consistent amino acid changes were observed in the viral RNAi suppressor gene 1A, nor elsewhere in the genome in any of the host backgrounds. Our study indicates that the RNAi response restricts the sequence space that can be explored by viral populations. Moreover, our study illustrates how evolution towards higher virulence can be a highly reproducible, yet unpredictable process.

## Introduction

Viruses are ubiquitous parasites of cellular life and the most abundant biological entities on earth, predicted to appear in any replicator system (1–3). Virus-host coevolution is often understood as an arms race between host defense and viral counter-defense mechanisms that occurs in all host systems (4). For example, the interferon response, one of the most potent antiviral mechanisms in mammals, can be antagonized at the transcriptional, translational, or functional level by many different viruses (5).

In insects, RNA interference (RNAi) plays a critical role in antiviral defense, as viral double stranded RNA (dsRNA) is processed by the endonuclease Dicer-2 (Dcr-2) into small interfering RNAs (siRNAs), serving as guides for the endonuclease Argonaute-2 (AGO2) to mediate viral RNA degradation (6). Several insect viruses have been shown to suppress this pathway, illustrating how immune pressure exerted by RNAi affects the genetic makeup of a virus (7, 8). For instance, the Drosophila C virus (DCV) 1A protein binds double-stranded RNA (dsRNA), interacts with Dcr-2, and inhibits Dcr-2 nuclease activity (9,10), whereas proteins from two distinct RNA viruses, the cricket paralysis virus 1A protein and the nora virus VP1 protein bind and inhibit AGO2 function (10–13).

Upon infection, RNA viruses create swarms of closely related within-host genotypes, often referred to as viral quasispecies (14), that contribute to the adaptability of a viral population to changing environments (15, 16). They are the result of the high mutation rate of the viral RNA dependent RNA polymerase (RdRP) and the large population sizes of RNA viruses, which can be as high as 10^12^ infectious particles in an infected host (17, 18). Most mutations in this diverse virus population will be deleterious, some neutral, and very few beneficial (19, 20), and the frequency of these variants will fluctuate according to evolutionary forces, such as the positive and purifying selection imposed by the host immune system. New virus variants that are capable of escaping immune surveillance have an evolutionary advantage and will likely take over the ancestral population. In accordance, plants with a more restrictive immune system have been shown to drive faster evolution of turnip mosaic virus (21).

In this study, we assessed to what extent selection pressure by the immune system drives virus population evolution. We used as a model system the fruit fly *Drosophila melanogaster* infected with its natural pathogen DCV, a single-stranded positive-sense RNA virus of the *Dicistroviridae* family (genus *Cripavirus*). The virus was serially passaged in wild-type flies and in fly mutants in which *Dcr-2* was either overexpressed or inactivated. Afterwards, the evolved virus populations were characterized using next-generation sequencing (NGS) in combination with phenotypic assays for virus replication and host survival. We found that the increase in the number of polymorphisms during passaging was most pronounced in *Dicer-2* knockout flies. However, the RNAi response had little effect on the accumulation of specific amino acid changes in viral proteins. Instead, we observed parallel evolution of mutations in the VP3 capsid protein in all viral lineages, which markedly enhanced viral replication and virulence *in vivo*.

## Results

### Experimental DCV evolution increases virulence

To investigate the effect of the host immune system on DCV evolution, we performed an experimental evolution study in which a parental DCV stock was serially passaged over three fly genotypes with different RNAi activity (Fig. 1A). Specifically, we used wild-type flies (*w^1118^*, hereafter WT), flies with a frameshift mutation in *Dcr-2* resulting in a premature stop codon (*Dcr-2^L811fsX^* (22), hereafter *Dcr-2-KO*), and flies overexpressing a *Dcr-2* transgene controlled by the UAS upstream activation sequence and a *Tubulin-Gal4* driver (hereafter *Dcr-2-OE*). As expected (23, 24), *Dcr-2* expression correlated with survival after DCV infection, with *Dcr-2-KO* flies having the shortest life span and *Dcr-2-OE* flies the longest (Fig. 1B).

**Figure 1.**
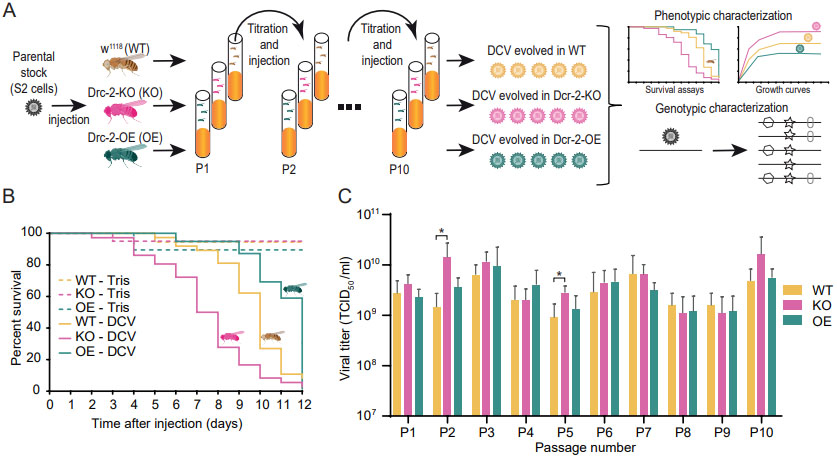
Experimental evolution of DCV in flies differing in their RNAi response. (**A**) A parental stock of DCV Charolles strain was serially passaged in wild-type flies (*w^1118^*; WT), *Dicer-2* knockout flies (*Dcr-2-KO*), and flies overexpressing *Dicer-2* (*Dcr-2-OE*) for ten fly generations with five independent lineages per genotype. Flies were inoculated by intrathoracic injections and titers were determined after every passage. Evolved virus populations were characterized phenotypically using growth curves and survival assays and genotypically by next-generation sequencing. (**B**) Survival curves of WT, *Dcr-2-KO* and *Dcr-2-OE* flies inoculated with 1,000 TCID_50_ of DCV. Two to five-day-old female WT flies were inoculated intrathoracically (n = 30) with the parental stock and survival was monitored daily. (**C**) Viral titers of all lineages at each viral passage. TCID_50_, median tissue culture infectious dose. Bars represent means and SD of the five viral lineages. Asterisks indicate *p* < 0.05, t-test. Fly and virus icons were retrieved from BioRender.

For each host genotype, we allowed virus populations to independently evolve during ten serial passages, with five independent lineages per genotype. The parental DCV stock was cultured in *Drosophila* S2 cells, and its genetic diversity had been reduced by three rounds of serial dilution prior to the experimental evolution study. To prevent population bottlenecks, flies were infected by intrathoracic injection with a high viral inoculum of 10^4^ median tissue culture infectious doses (TCID_50_), and virus titers were measured after each passage to normalize the inoculation of naive flies for the next passage (Fig. 1C). With the exception of passage 3 and 5, where we observed significantly higher viral titers in *Dcr-2-KO* than in WT flies (*p* < 0.05, *t*-test, Fig. 1C), the differences in viral titers between genotypes were generally small and not consistent along passages, likely because flies were harvested at the plateau of peak titers.

To examine the virulence of the evolved virus populations, we performed survival assays after inoculation of WT flies with 10^4^ TCID_50_ of the parental stock and virus stocks harvested after passage 1, 5 and 10 (P1, P5, P10; Fig. 2A-D). P10 virus populations from all lineages of all host genotypes induced higher mortality in WT flies than the parental stock (Log-rank test, *p* < 0.0001; hazard ratios > 3, Supplementary Table S1), with mean survival times decreasing from 5.5 days for the parental stock to 3.4 days for the P10 virus populations (average of all genotypes and lineages). The mean survival time of WT flies decreased from P1 to P5 for virus populations from all genotypes (paired *t*-test, *p* < 0.005) and further decreased from P5 to P10 only for virus populations from WT flies (paired *t*-test, *p* = 0.006) (Fig. 2D).

**Figure 2.**
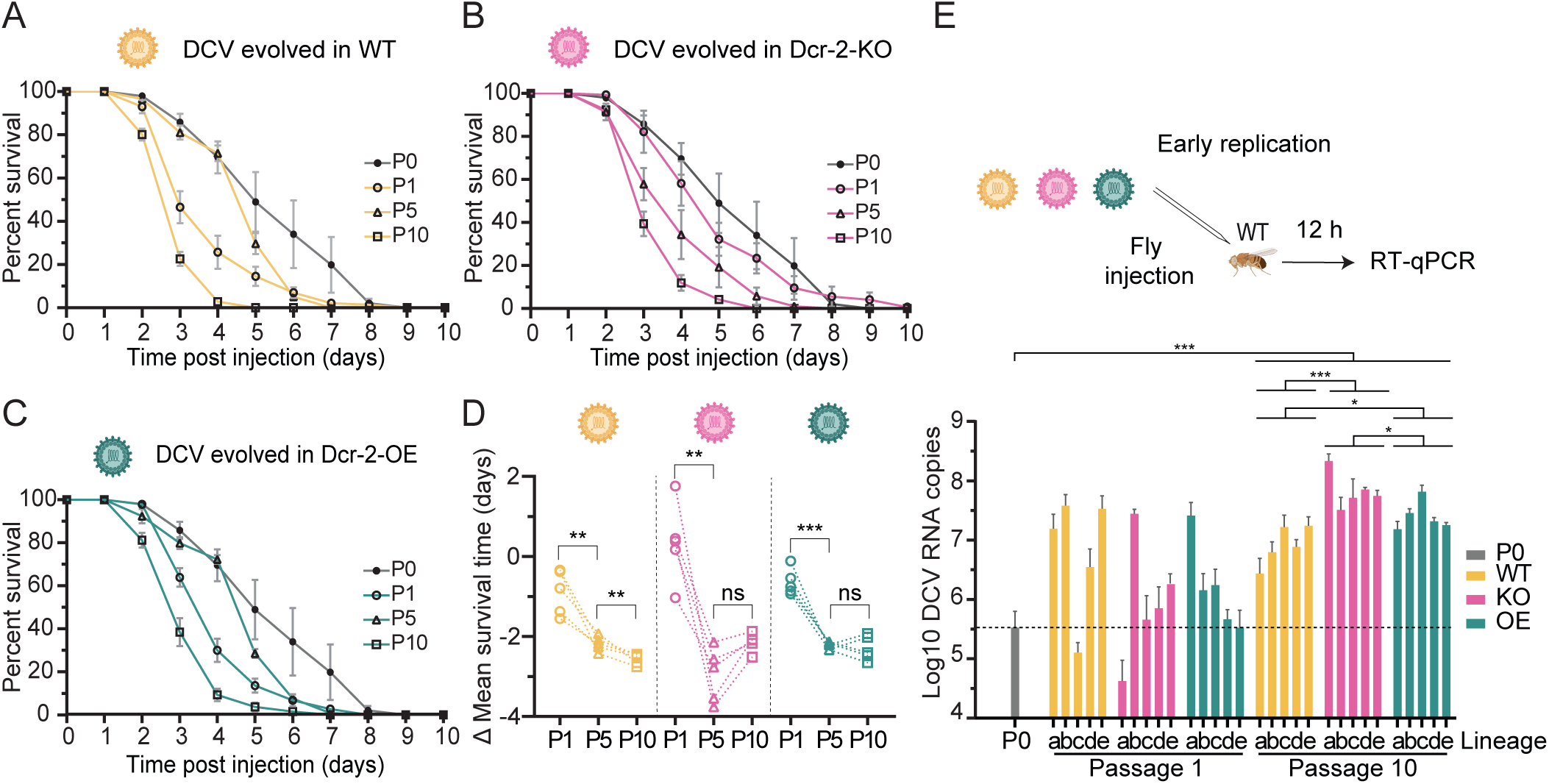
Increased virulence of evolved virus populations independent of the host genotype. (**A**–**C**) Survival curves of WT flies inoculated with 10,000 TCID_50_ of the parental DCV stock (P0) and virus populations from the indicated host genotypes at passage 1, 5 or 10. Two to five-day-old female WT flies (n = 30) were inoculated intrathoracically and survival was monitored daily. Results are shown as means and SEM of all five lineages for virus populations from (A) WT, (B) *Dcr-2-OE*, and (C) *Dcr-2-KO* flies. Data for the parental stock are shown as means and SEM of four independent experiments. Results from Cox regressions are shown in Supplementary Table S1. (**D**) Difference of the mean survival time for each virus stock relative to the parental stock. Dashed lines connect the values of each lineage. Asterisks indicate significance: **, *p* < 0.01; ***, *p* < 0.001, paired *t*-test. (**E**) Early replication kinetics of parental and evolved virus populations. 1,000 TCID_50_ of each virus stock were injected into WT flies and three groups of five flies were collected at 12 hours post infection (hpi). Viral RNA copy numbers were quantified by RT-qPCR. The dashed line shows the RNA copies of the parental stock. Results are shown as means and SD of three replicates. Viral RNA levels of the P10 virus populations and the parental stock were compared using a nested ANOVA followed by Tukey’s HSD test for multiple comparison. Adjusted *p*-values of Tukey’s HSD test are indicated with asterisks: *\*, p <* 0.05; ***, *p* < 0.001.

We next analyzed whether these survival phenotypes correlated with changes in replication kinetics of the evolved virus populations. To test this, we inoculated WT flies with the parental, P1, and P10 virus stocks and determined viral RNA levels at 12 hours post infection (hpi) (Fig. 2E). We observed a large variation in viral RNA replication between the independent lineages of all genotypes at P1, with the parental stock accumulating 3.4 x 10^5^ RNA copies and the P1 viruses ranging from 4.2 x 10^4^ to 3.9 x 10^7^ RNA copies. Irrespective of the genotype, all lineages of the P10 virus populations reached higher viral RNA levels (range 2.8 x 10^6^ to 2.2 x 10^8^ RNA copies) than the parental stock (nested ANOVA with significant effect of genotypes on the viral RNA level: *p* < 0.001, followed by Tukey’s HSD test with adjusted *p*-value: *p*-adj. < 0.0001). This phenotype was maintained when the infection was performed orally (Supplementary Fig. S1), indicating that our experimental evolution scheme did not lead to loss of infectivity via the presumed natural infection route. Among genotypes, P10 virus populations from *Dcr-2*-*KO* flies reached the highest RNA loads, compared to those evolved in WT and *Dcr-2*-*OE* flies (*p*-adj. = 0.0003 and *p*-adj. = 0.0468 Tukey’s HSD test), where those evolved in *Dcr2*-*OE* flies reached higher RNA loads compared to WT (*p*-adj. = 0.0374 Tukey’s HSD test) (Fig. 2E). Together, these results indicate that *in vivo* serial passage increases replication kinetics and virulence of the evolved virus populations.

### Ongoing accumulation of polymorphisms

We next investigated how the virus populations evolved through the serial passage experiment by next-generation sequencing (NGS). Viral genomes were sequenced to high coverage (average coverage of 127,542, Supplementary Fig. S2), allowing us to confidently call low-frequency mutations, as the median Phred sequencing quality score for the mutation calls was 39, corresponding to an error rate of 0.0001. We observed increasing numbers of mutations during passaging with a mean of 381 polymorphic sites in the evolved virus populations after ten passages, 116 more than in the parental stock (Fig. 3A). Between 179 and 499 single nucleotide variants (SNVs) per evolved population occurred at a frequency below 1%, and between 5 and 21 SNVs at a frequency higher than 1% (Fig. 3B). The number of polymorphic sites increased significantly through the serial passage in *Dcr-2-KO* and *Dcr-2-OE* flies (*p* < 0.05, paired *t*-test, Fig. 3A). This increase was most pronounced in virus populations from *Dcr-2-KO* flies, which accumulated a mean of 444 polymorphic sites at passage 10 (Fig. 3A), which was significantly different from virus populations from WT and *Dcr-2-OE* flies (*p* < 0.02, *t*-test). Yet, the vast majority of polymorphic sites were low-frequency mutations (median: 0.1%), and very few SNVs increased in frequency or reached fixation (Fig. 3B).

**Figure 3.**
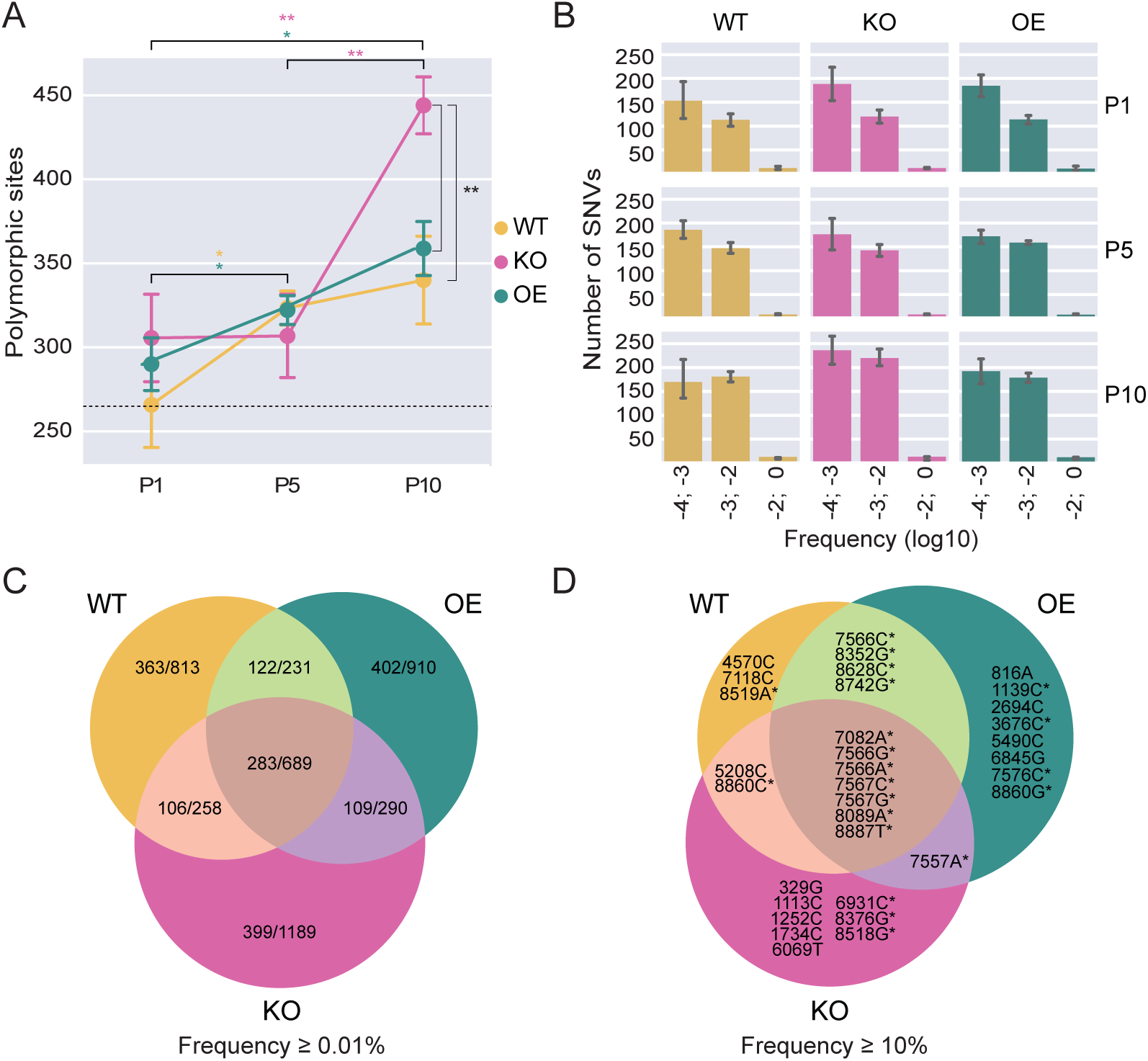
Increasing genetic variation during serial passage. **(A)** Number of polymorphic sites of the virus populations from WT, *Dicer-2-OE* and *Dicer-2-KO* flies across passages. Data are shown as means and SEM across the five lineages for each genotype. The dashed line marks the number of polymorphic sites in the parental stock with respect to the Charolles reference sequence (GenBank Accession Number: MK645242.1). **(B)** Frequencies of single-nucleotide variants (SNVs) in the indicated bins for virus populations from the indicated genotypes. Error bars indicate 95% confidence intervals based on the bootstrap distribution. **(C**–**D)** Venn diagram of the number of unique and shared SNVs between virus populations from WT, *Dicer-2-OE* and *Dicer-2-KO* flies. Per genotype, SNVs from all lineages and passages with a minimum observed frequency of 0.01% (C) or 10% (D) were combined. The ratios in (C) indicate the number of non-synonymous mutations to all mutations. Asterisks in (D) indicate non-synonymous mutations.

We next analyzed virus population diversity over time and found that the mean nucleotide diversity did not change significantly over time in any of the host genotypes, nor were there significant differences in population diversity between genotypes (*p* > 0.05, mixed ANOVA, Supplementary Fig. S3A). Mean nucleotide diversity was significantly higher in coding than in non-coding regions of the viral genome (*p* < 0.01, mixed ANOVA, Supplementary Fig. S3A). Specifically, the gene encoding the viral VP3 protein displayed a significant increase in nucleotide diversity compared to the mean diversity of the whole coding region (*p* < 0.05, paired *t*-test, Supplementary Fig. S3B).

### Moderate host genotype-specific adaptation in *Dicer-2* mutants

We next assessed the shared and unique SNVs across genotypes, first considering SNVs with frequencies ≥ 0.01%, and found a similar number of unique SNVs between virus populations from WT and *Dcr-2-OE* flies (n = 813 and 910, respectively), both having less SNVs than virus populations from *Dcr-2-KO* flies (n = 1189) (Fig. 3C). 15.7% of the SNVs were shared between virus populations from all three fly genotypes, of which 41.07% were non-synonymous mutations.

When restricting the analysis to high-frequency SNVs (≥ 10%), the fraction of SNVs shared by the virus populations from all three fly genotypes increased to 21.3%, all of which were non-synonymous mutations (Fig. 3D).

To understand if the observed mutation patterns are the result of host-specific adaptation, we conducted a permutation test, as described by Bons et al. (25). This test assesses whether SNVs occur at higher rates in specific genotypes than expected under a null hypothesis that SNVs are equally likely to occur in each genotype. For virus populations from WT and *Dcr-2-OE* flies, we did not find significant differences between the expected and observed number of SNVs. In contrast, 131 SNVs were unique to virus populations from *Dcr-2-KO* flies, which was significantly different from the expected number by random allocation (n = 89; *p* < 0.001, Supplementary Fig. S4). Together, these data indicate that viral populations are predominantly shaped by host-genotype independent, parallel evolution of shared SNVs as well as moderate host-genotype specific adaptation in the absence of a functional RNAi response.

### Deletions in homopolymeric uridine tracts

To explore the evolution of the viral populations in more detail, we visualized the SNVs across the DCV genome in a heatmap (Fig. 4A). We observed some SNVs that were already present in the parental stock and were retained in the viral populations throughout the experiment in most of the lineages (nt 329, 504, 2686, 2736, 7082, 7458, 8089 and 8526), while others appeared *de novo* during the experiment in different lineages (e.g., nt 331, 1026, 4758, 6263 and 7557). Strikingly, we found a ubiquitous deletion of a uridine in a poly(uridine) tract at position 5765– 5774, which gives rise to a frameshift and a premature stop codon in the *RdRP* gene. This deletion was also present in the parental stock and it was retained at low, yet fairly constant frequency throughout the experiment in all 15 lineages (around 4%, Supplementary Table S2). Likewise, another uridine deletion in a tract of 8 uridines at position 276–284 in the 5’ untranslated region (UTR) occurred in all 15 lineages with a constant frequency of around 3% (Supplementary Table S2). To control for possible sequencing artifacts, we evaluated if the two uridine deletions were located at the edges of the sequence reads containing them, as there is lower confidence in the base calling at read edges. However, in only 7% of reads the deletions were located in proximity to the read edges (within five nucleotides). Furthermore, the mean sequencing quality score in the uridine tracks and the adjacent positions was above 40 both in reads with and without deletion, corresponding to an error rate of 0.0001 (mean quality scores across uridine tracks: 40.1; 10%-quantile: 38.2, 90%-quantile: 41.0; Supplementary Table S3). Further analysis suggested that such deletions also occur in other, published DCV NGS datasets, and we suggest that the viral RdRP is prone to slippage on homopolymeric uridine tracts and that there is purifying selection against the occurrence of these tracts in RNA viruses (Supplementary text S1, Supplementary Table S4-S6).

**Figure 4.**
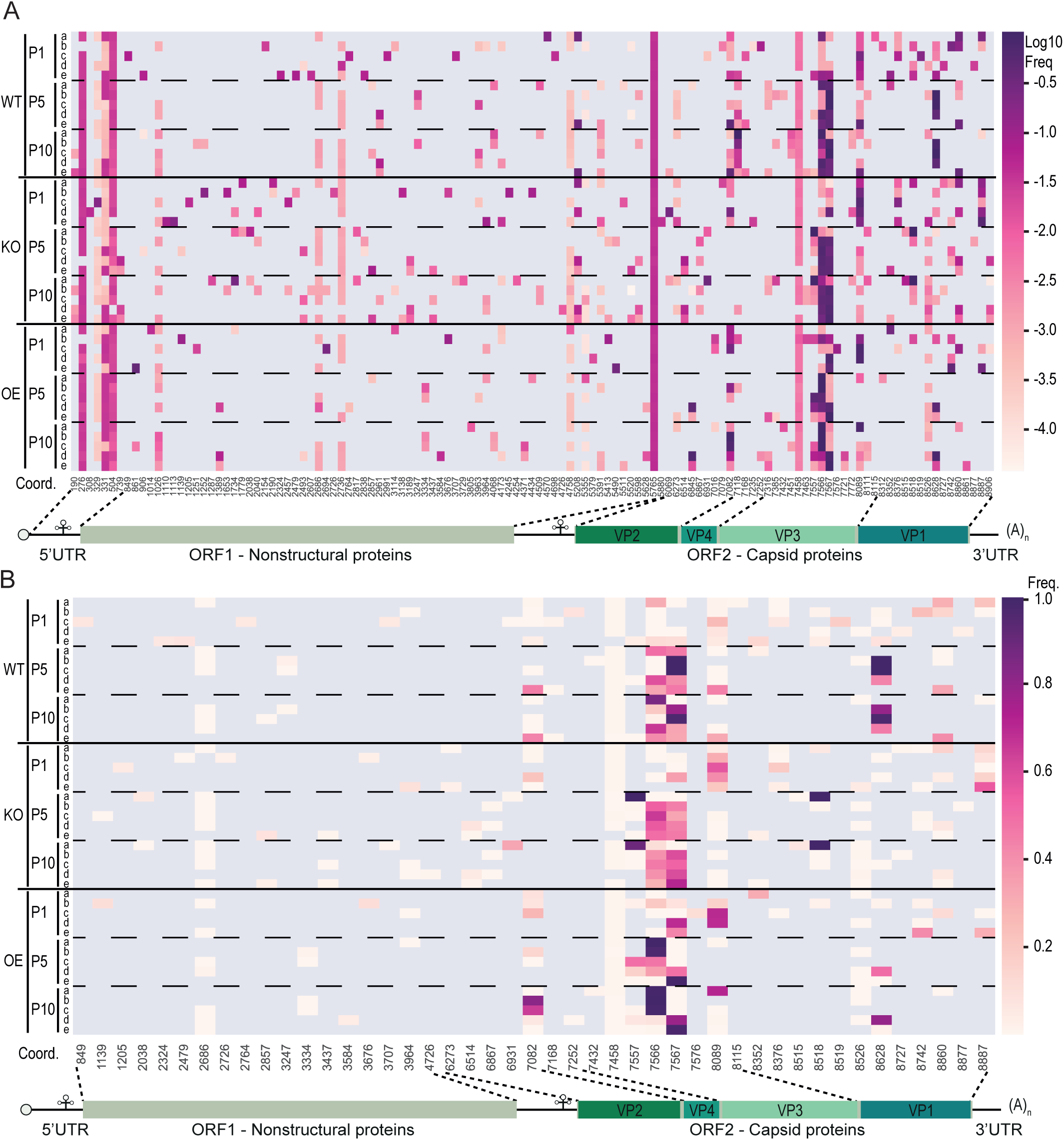
Parallel evolution of virus lineages. Heatmap of SNV frequencies of all SNVs **(A)** or non-synonymous SNVs **(B)** for virus populations from WT, *Dicer-2-OE* and *Dicer-2-KO* flies. The x-axis represents genome positions relative to the DCV EB reference strain (NC_001834.1). Only SNVs that occurred in at least one sample with an observed frequency ≥ 0.01 are shown. Deletions were excluded from the analysis in panel (B). A schematic representation of the DCV genome is shown below the heatmaps.

### Parallel evolution is driven by positive selection on the capsid protein

Thirteen SNVs became dominant (frequency > 50%) after ten passages, most of which changed the amino acid sequence of the capsid proteins in a host genotype-independent manner (Fig. 3D and Fig. 4B). Of note, two high-frequency SNVs appeared to be host genotype specific: 7118C became dominant only in WT flies (at a frequency of 99% in one lineage and above 20% in two others) and 6845G became dominant in two lineages from *Dcr-2-OE* flies. However, the majority of the dominant SNVs were located at codon 95 of the capsid *VP3* gene (nt 7566–7567), where the tyrosine codon (Y95) in the parental stock was replaced in 14 out of 15 lineages (Fig. 5A). After one passage, the Y95 frequency already declined below 60% in three out of five lineages in *Dcr-2-OE* flies, whereas it remained present at frequencies above 90% in four and three lineages in *Dcr-2-KO* and WT flies, respectively. After five passages, Y95 was undetectable in 12 out of 15 lineages, and at P10, Y95 was only retained in lineage D from *Dcr-2-OE* flies (at a frequency of 6%) and in lineage A from *Dcr-2-KO* flies, which was the only lineage in which Y95 remained dominant. Interestingly, in the latter lineage, two other capsid mutations, VP3 P92T and VP1 N114S, were fixated instead.

**Figure 5.**
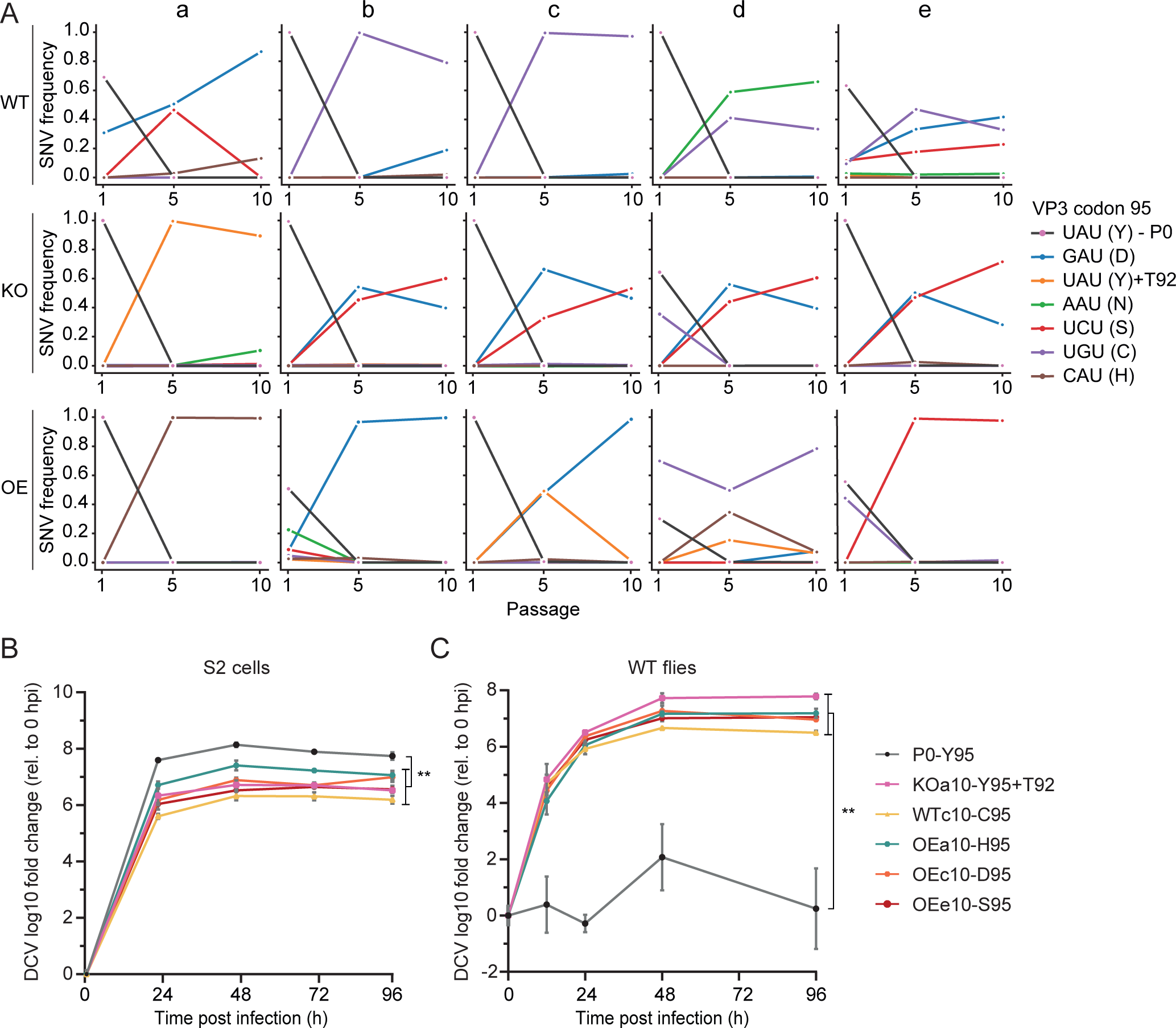
A capsid mutation increases *in vivo* viral fitness. **(A)** Frequencies of the codon coding for amino acid 95 of VP3 in DCV populations from the indicated host genotypes at passage 1, 5 and 10. UAU represents the parental codon coding for tyrosine (95Y). One variant retained the 95Y residue, but contained a substitution in its vicinity (P92T). **(B)** Growth curves of individual variants in *Drosophila* S2 cells infected at an MOI of 0.1. Variants were named according to the host genotype, lineage and passage number. P0 is the parental virus. Three wells were collected per time point for viral RNA quantification by RT-qPCR. Viral RNA levels are normalized to the 0 hpi time point and presented as geometric mean and SD. **(C)** Growth curves of individual variants in WT flies after inoculation with 100 TCID_50_ of each variant. Three pools of five flies were collected per time point for viral RNA quantification by RT-qPCR. Viral RNA levels are normalized to the 0 hpi time point and presented as geometric mean and SD. Differences in log10-transformed viral RNA levels between the parental and evolved lineages at 96 hpi were analyzed by *t*-tests (B) and Welch’s unequal variances *t*-tests (C). **, *p* < 0.01. Data for lineage OEa10 in (C) did not follow a normal distribution and a Mann-Whitney U test was performed instead *(p* = 0.05).

Strikingly, from all nine possible SNVs that can be formed from the parental UAU codon coding for Y95, five were found in the evolved virus populations, coding for aspartic acid, asparagine, cysteine, serine and histidine. The emergence of these variants did not correlate with the host genotype in which the virus populations had evolved. The remaining four possible codons were not detected, two of which being stop codons, one coding for a synonymous substitution, and one coding for phenylalanine, the amino acid most similar to tyrosine. Overall, these results suggest that the presence of a benzene ring at residue 95 of VP3 is detrimental to virus infection in flies. To confirm the selection forces statistically, we computed the population nucleotide diversity per synonymous (piS) and non-synonymous (piN) site for each viral gene separately (Supplementary Fig. S5). There were significant main effects of viral gene, site type (synonymous and non-synonymous) and passage number (*p* < 0.001, mixed ANOVA), and significant interactions between viral gene and site type (*p* < 0.0001) and between gene and passage number (*p* < 0.05), but no significant effects of host genotype on the population nucleotide diversity (*p* > 0.05). We found clear indication of positive selection (piN - piS > 0) in VP3 across all passages (P1: *p*-adj.

= 0.01; P5: *p*-adj. < 0.0001; P10: *p*-adj. = 0.004, Tukey’s HSD test with Bonferroni-adjusted *p*-values for each passage). These results indicate positive selection for amino acid changes in the capsid VP3 protein.

### An exposed amino acid in the capsid is responsible for environment specific fitness gains

We hypothesized that the VP3 mutations were responsible for the observed fitness gains of the passaged virus populations (Fig. 2E). To test this, we selected P10 lineages with one of the fixated mutations (frequency ≥ 90%), performed a serial dilution to remove minority variants, and analyzed growth kinetics in the Drosophila S2 cell line and in WT flies inoculated with a low dose (multiplicity of infection of 0.1 and 100 TCID_50_, respectively). All variants efficiently replicated in S2 cells, but the parental stock showed higher replication kinetics than the other variants (unpaired *t*-test, *p* < 0.01, Fig. 5B). In contrast, the parental stock scarcely replicated in adult flies at this low inoculum, whereas the evolved variants replicated efficiently *in vivo* (unpaired Welch’s *t*-test, *p* < 0.01, Fig. 5C). These results suggest that Y95 may be beneficial for DCV replication in cell culture, but detrimental for replication and virulence *in vivo*.

To investigate the effects of Y95 on the structural stability of the virion, we modeled the effect of the fixated mutations on the structure of the virus particle. We used the crystal structure of cricket paralysis virus (CrPV) (26), a related dicistrovirus which shares 60% sequence identity with the DCV capsid polyprotein and 79% identity within the region of interest. Given the high identity between these two proteins, it is possible to model the variants in CrPV capsid to predict their probable implications on the DCV capsid. The capsid polyprotein is a large assembly of 240 subunits, composed of copies of four proteins (VP1, VP2, VP3 and VP4) (Fig. 6A). The region of interest is located in a loop close to the protein-protein interface between VP2 and VP3 (Fig. 6A, 6C-6E). While this region is conserved across related dicistroviruses that share more than 50% sequence identity with the DCV capsid, the residues at positions 92 and 95 are relatively poorly conserved (Fig. 6B). Moreover, given their location in the loop, these residues are predicted to be solvent-exposed, suggesting that they may not be involved in maintaining the structural integrity of the capsid. Indeed, we do not find drastic differences in the predicted folding free energy of the capsid structure between the parental Y95 and its mutants (ranging from 0.71 kcal/mol to 3.33 kcal/mol; Supplement Table S7), suggesting that the observed fitness gains cannot be explained by structural effects of the acquired mutations.

**Figure 6.**
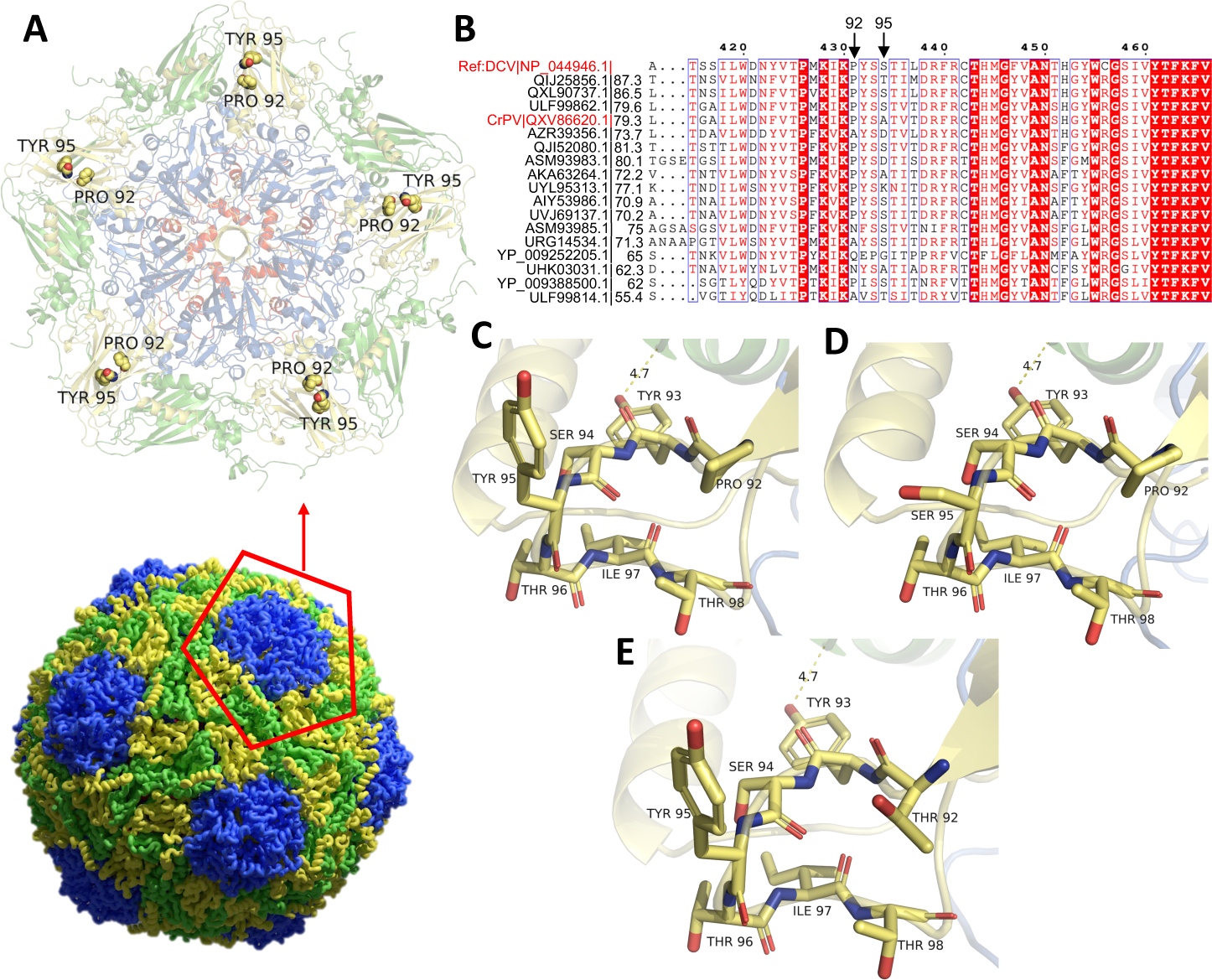
Structural analysis of capsid mutations selected during serial passage. **(A)** The CrPV capsid structure (PDB: 1B35) is shown with VP1 in blue, VP2 in green, VP3 in yellow, and VP4 in red (bottom). View at the fivefold symmetry axis of the capsid in cartoon representation with Tyr95 and Pro92 highlighted as spheres (top). **(B)** Multiple sequence alignment of the capsid of DCV and related viruses in the VP3 region of interest. Accession numbers are followed by their amino acid sequence identities to the DCV reference sequence (NP_044946.1). Completely conserved residues are highlighted in red blocks. **(C)** The parental residues Tyr95 and Pro92 are located in a solvent-exposed loop connecting two beta strands. Amino acids surrounding Tyr95 and Pro92 within a distance of 6Å are shown as sticks. **(D)** A similar view showing the Tyr95 to Ser95 substitution selected upon serial passage. **(E)** A marginally stabilizing mutant Pro92Thr (-1.79 kcal/mol) is shown. Proximity of the loop to VP2 (green) at a distance of 4.7Å is depicted in panels (C–E).

## Discussion

In this study, we have explored the effect of immune pressure on virus evolution using *Drosophila* as a model host. We passaged DCV in flies differing in Dcr-2 activity, hypothesizing that variations in RNAi immune pressure would affect virus evolution at the phenotypic and genotypic level, for instance by removing selection from the viral 1A protein that interferes with the host RNAi pathway. However, the effect of the RNAi machinery on DCV evolution was moderate. No specific polymorphisms accumulated in the DCV 1A gene in virus populations from *Dcr-2-KO* flies, nor in the other genotypes. However, we did observe that virus populations from *Dcr-2-KO* flies accumulated more polymorphisms than *Dcr-2* expressing flies, in line with previous observations (27), suggesting that the absence of immune pressure releases selective constraints, reducing purifying selection and allowing virus populations to further explore the mutational landscape.

It will be of interest to use our experimental approach to study the evolution of viruses that differ in their mode of RNAi suppression. Although DCV 1A interacts with several proteins including Dicer-2 (10), it relies on its dsRNA binding activity to suppress RNAi (9). As such, DCV may be under different selection pressure than viruses that suppress RNAi via direct protein-protein interactions, such as CrPV and Nora virus (10–13). It will likewise be interesting to study DCV evolution in flies bearing alleles of the *pastrel* restriction factor, which has strong effects on viral replication and virulence and may encode a protein that directly interacts with viral proteins (28).

Even though we observed some host genotype-specific effects on virus evolution, the strongest evolutionary pattern was shared among virus populations from all host genotypes, suggesting parallel and reproducible evolution. Specifically, the parental tyrosine at position 95 of VP3 was substituted with either one of five different amino acids, resulting in strongly enhanced replication and virulence *in vivo*. The predictability and reproducibility of evolutionary trajectories across independent populations has been a long-standing problem (29). Examples of parallel evolution exist, e.g., a mutation giving rise to antibiotic resistance was observed in multiple isolates of *Escherichia coli* independently (30), and specific drug resistance-associated mutations appeared repeatedly in human immunodeficiency virus-1 (HIV-1) in individuals receiving antiretroviral therapy (31–33). Likewise, the same subset of mutations has arisen independently in HIV-1 in two different cell lines during a long-term experimental evolution study (34). In contrast, in the long-term evolutionary study of Lenski *et al.* (35), the phenotype allowing *E. coli* to use citrate in aerobic conditions occurred in multiple independent lineages, but the underlying molecular mechanism was different (36). In our experiment, VP3 gene mutations occurred in parallel in all virus lineages, concomitant with a fitness gain and higher virulence *in vivo*. These mutations were fixated in 14 out of 15 lineages independently, indicating that loss of Y95 is a highly reproducible, yet unpredictable process, as five different variants have taken over the population.

Another instance of polymorphisms occurring in all lineages were deletions of a single uridine in homopolymeric tracts of at least 6 uridines in the 5’ UTR and in the RdRp gene. These uridine deletions also occurred in NGS data of DCV from other studies, and we propose that they are caused by RdRP slippage on homopolymeric uridine tracts. In contrast, thymidine deletions did not occur in NGS data from a DNA virus, herpes simplex virus 2, likely as a result of the proofreading activity of its viral DNA polymerase. In agreement, *E. coli* strains lacking mismatch repair mechanisms show increased numbers of deletions in simple sequences repeats (37).

It is well described that insertions and deletions preferentially occur in homopolymeric tracts in viral genomes (38). Some viruses make use of this feature, such as viruses of the *Potyviridae* family that use transcriptional slippage to insert an adenosine in a specific GAAAAAA sequence in viral transcripts to induce a frameshift and expression of an alternative viral protein (39). Likewise, the measles and Sendai viruses undergo a slippage event that adds an extra guanidine in transcripts coding for the P protein, creating a frameshift transcript that codes for the V protein involved in antagonizing host immunity (40). However, transcriptional slippage also occurs in picornaviruses on homopolymeric stretches of more than six uridines and these sequences are underrepresented in their genome, suggesting that there are also fitness costs associated with poly(uridine) tracts (41). The deletion in the *RdRP* gene in DCV introduces a premature stop codon in the RdRP and likely results in the production of defective viral genomes (DVGs). DVGs have been reported in many virus families and may affect pathogenesis via different mechanisms, like interfering with antiviral immunity or facilitating viral persistence (42). Whether DCV likewise produces DVGs and their involvement in the infection cycle remains to be established.

We found that the parental DCV stock replicated at low levels in flies receiving a low inoculum, but that it efficiently replicated in cell culture. This suggests that the parental virus is unfit because it lacks a basic function that is essential *in vivo*, but not in cell culture. Considering that the virus replicates efficiently in cells, we expect cell entry not to be impaired. Using structural models, we predicted that Y95 substitutions do not affect virus particle stability and that the residue is solvent exposed. We hypothesize that the observed phenotype is due to immune recognition of the parental stock containing Y95 in VP3, and escape thereof by mutants in VP3. This scenario is not unanticipated, as a single nucleotide substitution has been found to mediate immune evasion before. For example, blood clearance of alphaviruses like chikungunya virus is mediated mainly by resident macrophages in the liver, but a single amino acid substitution in the E2 glycoprotein disrupts this process and increases virus dissemination (43). We speculate that the parental virus is recognized by an immune receptor to mediate virus clearance, thus explaining the differences between *in vitro* versus *in vivo* replication.

To conclude, we have found that elimination of RNAi-based immune pressure during virus evolution led to a higher accumulation of polymorphisms, perhaps due to a reduction in purifying selection. Moreover, we have observed how a single amino acid change in the capsid of an RNA virus can have dramatic environment-specific effects on the growth rate. Our study illustrates how evolution towards higher virulence can be a highly reproducible, yet unpredictable process.

## Materials and Methods

### Fly strains and husbandry

Flies were maintained at 25 °C in standard fly food. Eggs were treated with bleach and, subsequently, flies were treated with tetracycline for two fly generations, as described before (44). Absence of nora virus, Drosophila C virus, Drosophila X virus and cricket paralysis virus was confirmed by RT-PCR using random hexamers for cDNA synthesis and the following primers for PCR:

NoraFor, ATGGCGCCAGTTAGTGCAGACCT;

NoraRev, CCTGTTGTTCCAGTTGGGTTCGA;

DCVFor, AAAATTTCGTTTTAGCCCAGAA;

DCVRev, TTGGTTGTACGTCAAAATCTGAG;

DXVFor, TCGGAAGAACCAAAAGGATG;

DXVRev, GTCCTCTCCACGCACTCTTC;

CrPVFor, ACGAGGAAGCAACTCAAGGA;

CrPVRev, GAGCCCGCTGAGATGTAAAG.

Absence of *Wolbachia* in the fly stocks was confirmed by PCR using primers wspFor TTTGCAAGTGAAACAGAAGG, wspRev GCTTTGCTGGCAAAATGG. Isogenized *Dicer-2-KO (Dcr-2^L811fsX^*) have been described before (27). Flies overexpressing *Dicer-2* were generated by crossing *UAS-Dicer-2* virgin females (VDRC ID 60008, (45)) with *Tubulin-Gal4*/*TM3 Sb* male flies, after which flies lacking the *Sb* marker were selected from the F1 offspring*. w^1118^* flies were used as wild-type. Two to five-day-old female flies were used for the serial passage and the RNA extractions.

### Virus stock and titration

The parental DCV stock (Charolles strain) was produced in *Drosophila* S2 cells, which were maintained in Schneider’s medium supplemented with 10% fetal calf serum and 50 U/ml penicillin and 50 μg/ml streptomycin (Gibco) medium at 27 °C. The stock was bottlenecked through three rounds of serial dilutions, by selecting the highest dilution that still showed cytopathic effects. Subsequently, a T25 flask of confluent S2 cells containing 5 ml of medium was infected with the bottlenecked virus. After 72 hours, the supernatant was collected, centrifuged at 300 *g* for 5 min, aliquoted, and stored at -80 °C. Titers were determined by end-point dilution in 96-well plates, as described before (44), and expressed as median tissue culture infectious dose (TCID_50_), calculated with the Reed-Muench method.

### Experimental evolution

Females of each genotype were intrathoracically inoculated with 10,000 TCID_50_ of DCV in 50 nl of 10 mM Tris-HCl, pH 7.3 using a Nanoject II microinjector (Drummond Scientific). For each of the five lineages per genotype, forty flies were inoculated. For the first passage, flies were inoculated with the parental stock and from P1 onwards, all 15 lineages were kept independently. After each passage, flies were harvested for preparation of virus stocks. Pools of ten flies were homogenized in 220 μl of PBS, using 1 mm silica beads in a Precellys homogenizer two times for 10 seconds, centrifuged at 16,000 *g* for 10 min to discard fly debris, and the supernatant was transferred to fresh tube, aliquoted, stored at -80°C, and titrated. For RNA extraction and NGS library of the parental stock, 1 ml of RNA-Solv (Omega) was added to 100 μl of virus stock, while for the viral lineages, pools of five flies were homogenized in 1 ml of RNA-Solv (Omega) using 1 mm silica beads in a Precellys homogenizer.

### Virus infections and survival assays

To study virulence, 10,000 TCID_50_ of all lineages from P1, P5, P10 and the parental DCV stock were injected separately into 35 female *w^1118^* flies. Survival was checked daily and analyzed using the Kaplan-Meier estimator. Early replication was assessed after inoculation of two to five-day-old female *w^1118^* flies with 1000 TCID_50_ of all lineages of P1, P10 or the parental stock. Three pools of five flies per lineage were collected at 12 hpi and processed for RT-qPCR. For the oral infections, 30 female *w^1118^* flies were inoculated with 1000 TCID_50_ by intrathoracic injection, placed in tubes for three days, after which these flies were discarded and 30 naive female *w^1118^* flies were added to the contaminated vial. Three pools of five flies were collected at 48 hpi and processed for RT-qPCR.

To analyze replication of the different capsid variants, P10 virus lineages containing the variant of interest at a frequency ≥ 90% were bottlenecked by serial dilution. Supernatant from the last well showing cytopathic effect in an end-point dilution was used to grow stocks in a T25 flask, followed by titration and Sanger sequencing to confirm the presence of the mutation. 100 *w^1118^* female flies were inoculated with 100 TCID_50_ of each viral stock, and three pools of five flies per variant were collected at the given time points and processed for RT-qPCR.

### RT-qPCR

Total RNA was extracted, reverse transcribed using the TaqMan reverse transcription kit (Thermo Fisher) using oligo (dT) as a primer and quantitative PCR analysis was performed using the GoTaq qPCR SYBR master mix (Promega) on a LightCycler 480 instrument (Roche), according to the manufacturers’ recommendations. DCV was quantified using primers DCVFor (TCAAGAAAAGTTGCGTGGGT) and DCVRev (CAGAGCGTCCTTGGAGAGTG), and expression was normalized to expression of *Ribosomal protein 49*, amplified using primers Rp49For (ATGACCATCCGCCCAGCATAC) and Rp49Rev (CTGCATGAGCAGGACCTCCA). A standard curve of 10-fold dilutions of plasmid pAc5.1-V5-His-A (Invitrogen) containing the DCV ORF-2 sequence was used to convert Ct values to genome copy numbers.

### RNA extraction, cDNA synthesis and NGS library preparation

Total RNA was isolated using RNA-Solv (Omega Bio-tek), and reverse transcribed using Superscript IV (Thermo Fisher) and oligo (dT) as a primer, following the manufacturers’ instructions. Viral cDNA was PCR amplified using CloneAmp HiFi (TaKaRa) in four overlapping amplicons of about 2.2 kb, using the following primers:

For1, TGTACACACGGCTTTTAGGTAGA;

Rev1, GGAAAAGTGTTGCAAGAGCGA;

For2, TGTTCTTCGGGAAATGGGGA;

Rev2, TGCCTGTCCCACACGAATAG;

For3, GTTACGCGTGTCCTTTGACG;

Rev3, AACTTCCGTACCAACGCTCA;

For4, AAAGTTGCGTGGGTTTGTGG;

Rev4, CGTGTAAGCAGGGCAGATAGT).

Amplicons were purified with the NucleoSpin kit (MACHEREY-NAGEL), sheared in a Bioruptor Pico (Diagenode) by 12 cycles of 30 sec of sonication and 30 sec of cooling in 1.5 ml Bioruptor tubes (Diagenode), mixed in equimolar proportions, and used to prepare the library with the Next ultra II DNA library prep kit (E7645, NEB) using 1 μg of DNA per sample as input. Samples were multiplexed with barcodes (E7335, E7500, E7710, E7730, NEB), and the libraries were sequenced on an Illumina HiSeq 4000 sequencer as paired-end 100 base reads by the GenomEast platform, a member of the ’France Genomique’ consortium. Image analysis and base calling were performed using RTA version 2.7.7 and bcl2fastq version 2.20.0.422.

### NGS data processing

NGS data were analyzed using a custom computational workflow (Supplementary Fig. S6). For processing the raw read data, the bioinformatics pipeline V-pipe (46) was integrated into the workflow. BWA-MEM (47) was used to align reads from the parental stock to the DCV EB reference strain that contains complete UTR sequences (GenBank accession number: NC_001834.1). Bcftools was used to generate a consensus sequence from the parental sequence, which was then used to align the raw reads from the evolutionary samples from passage 1, 5, and 10 using BWA-MEM. Mutations with respect to the parental stock consensus sequence were called using ShoRAH (48) with default parameters (posterior threshold of 0.9, alpha 0.1 and shift 3). The posterior threshold was chosen conservative to ensure that only mutations of high confidence are included in the subsequent analyses. Moreover, the Strand Bias Filter (49) was applied to filter out further variation of low confidence. Phred sequencing quality scores for mutation calls were accessed using pysamstats (https://github.com/alimanfoo/pysamstats). Further, to evaluate the differences of our parental stock to its ancestor, the Charolles strain, the parental stock sample was processed with respect to the Charolles reference sequence (GenBank Accession Number: MK645242.1). An adapted version of vcf_annotator (https://github.com/rpetit3/vcf-annotator) was used to annotate non-synonymous mutations. Population nucleotide diversity and number of polymorphisms were computed considering only mutations with a minimum frequency of 0.0001. SNPGenie’s within-pool analysis was used in the default settings and minimum frequency 0.0001 (50) to compute the population nucleotide diversity per synonymous (piS) and non-synonymous (piN) site.

### Permutation test for genotype specific adaptation

A permutation test as described before (51) was used to test for host genotype-specific adaptation. Mutations that reached the frequency threshold of 0.0001 in at least one evolutionary line were considered for this analysis. For each mutation, the number of lineages containing the mutation were counted. Then, each mutation occurrence was randomly reallocated to a lineage by using the number of mutations that occurred in each lineage as weights. In this manner, 1000 samples were drawn. The observed number of mutations that were unique or shared among genotypes was then compared to the number expected from the permutation samples, and a two-sided test was used to assess the difference. For each category, *p*-values were computed as 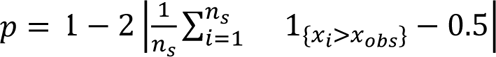 where 𝑛_s_ is the number of samples drawn, 𝑥_i_ is the 𝑖-th random realization and 𝑥_𝑎𝑏𝑠_ is the observed number of mutations in the respective category.

### Randomization test

A randomization test was performed to test whether the observed number of two consecutive phenylalanine (FF) residues encoded as ‘TTTTTT’ was different from the expected number based on random mixing of the TTT and TTC codons. For each viral genome, the total number of FF occurrences, the number of FF encoded by TTTTTT (𝑛_𝑇𝑇𝑇𝑇𝑇𝑇_), and the occurrences of the TTC and TTT codon were counted. For each viral genome, 𝑛_𝑠_ = 1000 samples were drawn, by randomly pairing the TTC and TTT codons based on their codon frequencies. To test whether the number TTTTTT encodings from the randomization exceeds the observed number, a two-sided test was performed. *P*-values were computed as 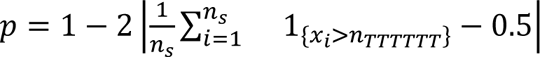 where 𝑥_𝑖_ denotes the number of TTTTTT realizations in the 𝑖-th randomization.

### Quality analysis of deletion calls

To evaluate the quality of reads supporting the two uridine deletions at positions 276–284 and 5765–5774, an additional quality control was performed on the alignment files of all samples. The mean position-wise base calling quality scores were assessed in the homopolymeric regions of the two deletions and the values were compared between reads with deletions to the values of reads without the deletions. In addition, the proportion of reads where the deletion track was located within a range of five nucleotides from the read edges was quantified.

### Capsid structural modeling

In the absence of structural information of the DCV capsid protein, the crystal structure of the capsid of CrPV (PDB: 1B35) was used to model the impact of mutations on capsid structure.

Mutations of interest, located at sites 92 and 95 in VP3, were modeled for all VP3 subunits in the capsid assembly. Structural models were constructed for each mutation using the functions RepairPDB and BuildModel from the FoldX suite (v.5.0) with five iterations of sidechain rotamer adjustments. For each mutant constructed, the average differences in capsid structural stabilities, i.e., free energy of protein folding (kcal/mol), between the parental amino acid and the mutants were calculated.

To assess sequence conservation of the DCV capsid polyprotein (NP_044946.1), a multiple sequence alignment was generated of closely related viruses (> 50% sequence identity), identified using a BLAST search: Warroolaba Creek virus (QIJ25856.1), Rondani’s wasp virus 1 (QXL90737.1), Cricket paralysis virus (QXV86620.1), Hypsignathus monstrous dicistrovirus (AZR39356.1), Millport beadlet anemone dicistro-like virus 1 (ASM93983.1), Maanshan Dicis tick virus 1 (UYL95313.1), Nilaparvata lugens C virus (AIY53986.1), Arma chinensis virus 1 (UVJ69137.1), Caledonia beadlet anemone dicistro-like virus 2 (ASM93985.1), Anopheles C virus (YP_009252205.1), Guiyang dicistrovirus 2 (UHK03031.1), Apis dicistrovirus (YP_009388500.1), as well as other, uncharacterized members of the *Cripavirus* genus (ULF99862.1) and the *Dicistroviridae* family (QJI52080.1, AKA63264.1, URG14534.1, ULF99814.1).

### Statistical tests

Two-sided *t*-tests on log10-transformed DCV titers for each passage was used for pairwise comparisons in viral titers in the different genotypes (Python package SciPy (52), ttest_ind function). Data were log10-transformed to ensure that the residuals were Gaussian distributed (Python package SciPy, normaltest p > 0.05 for each group), and equal variance was confirmed using a Levene’s test. Mean survival times of WT flies upon infection with virus populations from the different passages were compared across passages using a paired two-sided *t*-test. The Shapiro-Wilk test was applied to validate that the data were Gaussian distributed (Python package Pingouin (53), normality function). Mortality of WT flies infected with P10 viruses and the parental stock viruses was compared using a log-rank test and Cox regression (IBM SPSS Statistics, version 29.0.0.0). Viral RNA levels between the parental stock and the evolved viral populations at passage 10 were compared using a nested ANOVA, after which a Tukey’s HSD test with *p*-value adjustment was applied for multiple comparison between genotypes. The data were first log10-transformed, and equal variance was confirmed using a Levene’s test (*p* > 0.05). The factor ‘technical replicate’ is nested within each biological lineage, and the parental stock was classified as a separate genotype category. For each genotype, the difference in the number of polymorphic sites across passages was analyzed using paired two-sided *t*-tests. To compare differences in the number of polymorphic sites among the host genotypes at passage 10, two-sided *t*-tests were used (Python package SciPy, ttest_rel function). The data were Gaussian distributed (Python package, SciPy normaltest p > 0.05 for each group), and equal variance was confirmed using a Levene’s test. To analyze viral nucleotide diversity within passages and between fly genotypes, a mixed ANOVA was used with passages 1, 5 and 10 as within-subject factor and fly genotype as between-subject factor (Python package Pingouin, mixed_anova function). The residuals were Gaussian distributed, and equal variance was confirmed using a Levene’s test (Python package Pingouin, homoscedasticity function). To compare the mean nucleotide diversity between coding and non-coding regions, a mixed ANOVA was used with passages as the within-subject factor and coding or non-coding region as the between-subject factor (Python package Pingouin, mixed_anova function). A mixed ANOVA was performed to analyze the effect of genotype, passage, protein and site type (non-synonymous and synonymous) on the population nucleotide diversity. Passage was used as within-subject factor in the model. The data were cube-root-transformed to ensure approximate normal distribution and equal variance among groups, which was confirmed using a Levene’s test (*p* > 0.05). Subsequently, per passage, Tukey’s HSD test with Bonferroni adjusted *p*-value was used for multiple comparison of viral gene and site type. Viral RNA loads between different VP3 variants at 96 hpi in S2 cells were log-10 transformed and upon validation of the normal distribution using the Saphiro-Wilk test (function shaphiro.test, R Stats package) and equal variance by the F test (var.test function, R Stats package) and analyzed using two-sided *t*-tests. Welch’s unequal variances *t*-tests were used to compare log10-transformed viral RNA loads of VP3 variants at 96 hpi in WT flies. Unless noted otherwise, statistical analyses were performed using Rstudio 2022.02.1+461.

### Data availability

All raw NGS data have been deposited in NCBI BioProject under the accession number PRJNA993483.

### Code availability

The data processing pipeline can be found at https://github.com/cbg-ethz/DCV-Evolution-Data-Processing.

## Acknowledgements

We thank members of the laboratory for discussions and Carla Saleh (Pasteur Institute) for providing *Dicer-2* knockout flies. This work was financially supported by the European Union’s Horizon 2020 research and innovation program, under the Marie Skłodowska-Curie Actions Innovative Training Networks grant agreement no. 955974 (VIROINF).

## References

1. E. V. Koonin, V. V. Dolja, A virocentric perspective on the evolution of life. Curr. Opin. Virol. 3, 546–557 (2013).

2. J. Iranzo, P. Puigbò, A. E. Lobkovsky, Y. I. Wolf, E. V. Koonin, Inevitability of genetic parasites. Genome Biol. Evol. 8, 2856–2869 (2016).

3. M. Krupovic, V. V. Dolja, E. V. Koonin, Origin of viruses: primordial replicators recruiting capsids from hosts. Nat. Rev. Microbiol. 17, 449–458 (2019).

4. B. B. Finlay, G. McFadden, Anti-immunology: evasion of the host immune system by bacterial and viral pathogens. Cell 124, 767–782 (2006).

5. A. García-Sastre, C. A. Biron, Type 1 interferons and the virus-host relationship: a lesson in détente . Science 312, 879–882 (2006).

6. D. B. Gammon, C. C. Mello, RNA interference-mediated antiviral defense in insects. Curr. Opin. Insect Sci. 8, 111–120 (2015).

7. Z. Guo, Y. Li, S.-W. Ding, Small RNA-based antimicrobial immunity. Nat. Rev. Immunol. 19, 31–44 (2019).

8. A. W. Bronkhorst, R. P. van Rij, The long and short of antiviral defense: small RNA-based immunity in insects. Curr. Opin. Virol. 7, 19–28 (2014).

9. R. P. van Rij, et al., The RNA silencing endonuclease Argonaute 2 mediates specific antiviral immunity in *Drosophila melanogaster*. Genes Dev. 20, 2985–2995 (2006).

10. A. Nayak, et al., A viral protein restricts drosophila rnai immunity by regulating argonaute activity and stability. Cell Host Microbe 24, 542–557.e9 (2018).

11. A. Nayak, et al., Cricket paralysis virus antagonizes Argonaute 2 to modulate antiviral defense in Drosophila. Nat. Struct. Mol. Biol. 17, 547–554 (2010).

12. J. T. van Mierlo, et al., Novel Drosophila viruses encode host-specific suppressors of RNAi. PLoS Pathog. 10, e1004256 (2014).

13. J. T. van Mierlo, et al., Convergent evolution of argonaute-2 slicer antagonism in two distinct insect RNA viruses. PLoS Pathog. 8, e1002872 (2012).

14. A. S. Lauring, R. Andino, Quasispecies theory and the behavior of RNA viruses. PLoS Pathog. 6, e1001005 (2010).

15. S. Duffy, Why are RNA virus mutation rates so damn high? PLoS Biol. 16, e3000003 (2018).

16. A. S. Lauring, J. Frydman, R. Andino, The role of mutational robustness in RNA virus evolution. Nat. Rev. Microbiol. 11, 327–336 (2013).

17. S. Duffy, L. A. Shackelton, E. C. Holmes, Rates of evolutionary change in viruses: patterns and determinants. Nat. Rev. Genet. 9, 267–276 (2008).

18. A. Moya, E. C. Holmes, F. González-Candelas, The population genetics and evolutionary epidemiology of RNA viruses. Nat. Rev. Microbiol. 2, 279–288 (2004).

19. A. L. Hughes, Adaptive evolution of genes and genomes. Adaptive evolution of genes and genomes (1999).

20. A. Acevedo, L. Brodsky, R. Andino, Mutational and fitness landscapes of an RNA virus revealed through population sequencing. Nature 505, 686–690 (2014).

21. R. Navarro, et al., Defects in plant immunity modulate the rates and patterns of RNA virus evolution. Virus Evol. 8, veac059 (2022).

22. Y. S. Lee, et al., Distinct roles for Drosophila Dicer-1 and Dicer-2 in the siRNA/miRNA silencing pathways. Cell 117, 69–81 (2004).

23. D. Galiana-Arnoux, C. Dostert, A. Schneemann, J. A. Hoffmann, J.-L. Imler, Essential function *in vivo* for Dicer-2 in host defense against RNA viruses in *Drosophila*. Nat. Immunol. 7, 590–597 (2006).

24. M. J. Spellberg, M. T. Marr, FOXO regulates RNA interference in Drosophila and protects from RNA virus infection. Proc Natl Acad Sci USA 112, 14587–14592 (2015).

25. E. Bons, C. Leemann, K. J. Metzner, R. R. Regoes, Long-term experimental evolution of HIV-1 reveals effects of environment and mutational history. PLoS Biol. 18, e3001010 (2020).

26. J. Tate, et al., The crystal structure of cricket paralysis virus: the first view of a new virus family. Nat. Struct. Biol. 6, 765–774 (1999).

27. V. Mongelli, et al., Innate immune pathways act synergistically to constrain RNA virus evolution in Drosophila melanogaster. *Nat*. Ecol. Evol. 6, 565–578(2022).

28. M. M. Magwire, et al., Genome-wide association studies reveal a simple genetic basis of resistance to naturally coevolving viruses in Drosophila melanogaster. PLoS Genet. 8, e1003057 (2012).

29. S. J. Gould, Wonderful life: the Burgess Shale and the nature of history. Wonderful life: the Burgess Shale and the nature of history (1989).

30. J. P. M. Coolen, et al., Genome-wide analysis in Escherichia coli unravels a high level of genetic homoplasy associated with cefotaxime resistance. Microb. Genom. 7 (2021).

31. W. F. Flynn, et al., Deep sequencing of protease inhibitor resistant HIV patient isolates reveals patterns of correlated mutations in Gag and protease. PLoS Comput. Biol. 11, e1004249 (2015).

32. K. A. Crandall, C. R. Kelsey, H. Imamichi, H. C. Lane, N. P. Salzman, Parallel evolution of drug resistance in HIV: failure of nonsynonymous/synonymous substitution rate ratio to detect selection. Mol. Biol. Evol. 16, 372–382 (1999).

33. N. Beerenwinkel, et al., Estimating HIV evolutionary pathways and the genetic barrier to drug resistance. J. Infect. Dis. 191, 1953–1960 (2005).

34. F. Bertels, C. Leemann, K. J. Metzner, R. Regoes, Parallel evolution of HIV-1 in a long-term experiment. Mol. Biol. Evol. 36, 2400–2414. (2019).

35. R. E. Lenski, M. R. Rose, S. C. Simpson, S. C. Tadler, Long-Term Experimental Evolution in Escherichia coli. I. Adaptation and Divergence During 2,000 Generations. Am. Nat. 138, 1315 (1991).

36. Z. D. Blount, C. Z. Borland, R. E. Lenski, Historical contingency and the evolution of a key innovation in an experimental population of Escherichia coli. Proc Natl Acad Sci USA 105, 7899–7906 (2008).

37. P. Kumar, H. A. Nagarajaram, A study on mutational dynamics of simple sequence repeats in relation to mismatch repair system in prokaryotic genomes. J. Mol. Evol. 74, 127–139 (2012).

38. E. Domingo, “Molecular basis of genetic variation of viruses” in Virus as Populations, (Elsevier, 2020), pp. 35–71.

39. A. Olspert, B. Y.-W. Chung, J. F. Atkins, J. P. Carr, A. E. Firth, Transcriptional slippage in the positive-sense RNA virus family Potyviridae. EMBO Rep. 16, 995–1004 (2015).

40. J. F. Atkins, G. Loughran, P. R. Bhatt, A. E. Firth, P. V. Baranov, Ribosomal frameshifting and transcriptional slippage: From genetic steganography and cryptography to adventitious use. Nucleic Acids Res. 44, 7007–7078 (2016).

41. H. Stewart, A. Olspert, B. G. Butt, A. E. Firth, Propensity of a picornavirus polymerase to slip on potyvirus-derived transcriptional slippage sites. J. Gen. Virol. 100, 199–205 (2019).

42. M. Vignuzzi, C. B. López, Defective viral genomes are key drivers of the virus-host interaction. Nat. Microbiol. 4, 1075–1087 (2019).

43. K. S. Carpentier, et al., Discrete viral E2 lysine residues and scavenger receptor MARCO are required for clearance of circulating alphaviruses. eLife 8 (2019).

44. S. H. Merkling, R. P. van Rij, Analysis of resistance and tolerance to virus infection in Drosophila. Nat. Protoc. 10, 1084–1097 (2015).

45. G. Dietzl, et al., A genome-wide transgenic RNAi library for conditional gene inactivation in Drosophila. Nature 448, 151–156 (2007).

46. S. Posada-Céspedes, et al., V-pipe: a computational pipeline for assessing viral genetic diversity from high-throughput data. Bioinformatics 37, 1673–1680 (2021).

47. H. Li, R. Durbin, Fast and accurate short read alignment with Burrows-Wheeler transform. Bioinformatics 25, 1754–1760 (2009).

48. O. Zagordi, A. Bhattacharya, N. Eriksson, N. Beerenwinkel, ShoRAH: estimating the genetic diversity of a mixed sample from next-generation sequencing data. BMC Bioinformatics 12, 119 (2011).

49. K. McElroy, O. Zagordi, R. Bull, F. Luciani, N. Beerenwinkel, Accurate single nucleotide variant detection in viral populations by combining probabilistic clustering with a statistical test of strand bias. BMC Genomics 14, 501 (2013).

50. C. W. Nelson, L. H. Moncla, A. L. Hughes, SNPGenie: estimating evolutionary parameters to detect natural selection using pooled next-generation sequencing data. Bioinformatics 31, 3709–3711 (2015).

51. E. Bons, F. Bertels, R. R. Regoes, Estimating the mutational fitness effects distribution during early HIV infection. Virus Evol. 4, vey029 (2018).

52. P. Virtanen, et al., SciPy 1.0: fundamental algorithms for scientific computing in Python. Nat. Methods 17, 261–272 (2020).

53. R. Vallat, Pingouin: statistics in Python. JOSS 3, 1026 (2018).

