## Supplementary Text, Fig. S1 to S6, Table S1, S4, S5, S7 for "Parallel evolution and enhanced virulence upon *in vivo* passage of an RNA virus in *Drosophila melanogaster*"

**This PDF file includes:**

Supporting text
Tables S1 to S7
Figures S1 to S6
SI References

### Supplemental text

#### Uridine deletions in homopolymeric tracts

Single nucleotide deletions were observed at two locations in the DCV genome at low, yet fairly constant levels during passaging. Since both deletions occurred in homopolymeric uridine tracts, these may be due to error-prone RNA replication by the viral RdRP or to an artifact of library preparation and sequencing. To assess this, we first called variants in other, published DCV NGS data sets (1–3). Uridine deletions in those data sets occurred at the same positions as in our data, and at similar frequencies in the RdRP, but at variable frequencies in the 5'UTR (ranging from 3.6 to 87.8%) (Supplementary Table 3), suggesting that the deletion in the 5'UTR does not negatively impact virus fitness. We next called variants in Illumina based NGS data in a published dataset of a DNA virus, herpes simplex virus 2 (HSV2) (4). The HSV2 genome contains 38 tracts with more than six consecutive thymidines, but no deletions could be detected in those regions (Supplementary Table 5), suggesting that the occurrence of uridine deletions in our DCV data are not due to an sequencing artifact.

If uridine deletions are indeed due to error-prone RNA replication, we would expect negative selection against the occurrence of poly(U) tracts in viral genomes. We examined the coding regions of different RNA viruses for two consecutive phenylalanine (F) residues, which are encoded by either UUC or UUU and may cause the appearance of a hexameric uridine tract. With a randomization test, we then computed the expected number of FF to be encoded by UUUUUU, taking codon frequencies into account. For eight out of nine RNA viruses from different taxa analyzed (*Picornavirales, Mononegavirales*, and *Nidovirales* order), the observed number of UUUUUU was lower than the expected number, in three of these cases the underrepresentation reached statistical significance ($p < 0.05$; Supplementary Table 4). In contrast, for DNA viruses the expected number of poly-thymidine tracts matched the observed number for four out of six analyzed viruses (from the order *Herpesvirales, Lefavirales* and *Pimascovirales*), while the other two show a mild underrepresentation. Together, we suggest that the viral RdRP is prone to slippage on homopolymeric uridine and that there is purifying selection against the occurrence of these tracts in RNA viruses.

### Supplemental Tables

**Table S1.** Hazard ratios (RH) of the evolved virus population compared to the parental stock.

| | WT | | | | KO | | | | OE | | | |
|---|---|---|---|---|---|---|---|---|---|---|---|---|
| | | 95.0% CI | | | | 95.0% CI | | | | 95.0% CI | | |
| | RH | Lower | Upper | *p* value | RH | Lower | Upper | *p* value | RH | Lower | Upper | *p* value |
| **P1** | 2,12 | 1,63 | 2,75 | 1,76E-08 | 1,15 | 0,89 | 1,49 | 2,79E-01 | 1,87 | 1,44 | 2,42 | 2,56E-06 |
| **P5** | 1,63 | 1,23 | 2,16 | 5,80E-04 | 2,37 | 1,77 | 3,19 | 8,53E-09 | 1,59 | 1,19 | 2,12 | 1,55E-03 |
| **P10** | 4,37 | 3,21 | 5,94 | 7,45E-21 | 3,13 | 2,32 | 4,22 | 7,73E-14 | 3,38 | 2,51 | 4,55 | 8,30E-16 |

**Table S2.** Table of all SNVs with minimum frequency 0.0001 per evolutionary lineage. See separate file.

**Table S3.** Detailed quality control for deletion calls in uridine tracts at position 276-285 and position 5765-5774. See separate file.

**Table S4.** Frequencies of uridine deletions at position 276-285 and position 5765-5774 in DCV sequencing data from the literature. Data were analyzed using the same data processing pipeline as described in 'Methods'.

| Reference | Sample Accession Number | Position 276-285 | | Position 5765-5774 | |
|---|---|---|---|---|---|
| | | Coverage | Frequency | Coverage | Frequency |
| (1) | SRR17044453 | 18182 | 15.34% | 10108 | 2.15% |
| (1) | SRR6156525 | 1304 | 0.00% | 2469 | 3.81% |
| (2) | ERR3510613 | 805 | 15.78% | 1126 | 2.13% |
| (2) | ERR3510616 | 589 | 3.57% | 1310 | 0.00% |
| (3) | SRR17044471 | 29353 | 41.74% | 16762 | 2.2% |
| (3) | SRR17044453 | 14470 | 16.48% | 7933 | 2.28% |
| (3) | SRR17044623 | 19169 | 47.43% | 6067 | 2.46% |
| (3) | SRR17044490 | 24236 | 87.81% | 10541 | 3.19% |

**Table S5.** Underrepresentation of FF amino acid encoded by UUUUUU in viral genomes.

| Virus | Reference genome | Virus type | Number of FF | Observed number of UUUUUU encoding FF | Expected number of UUUUUU encoding FF, based on randomization | | | *p*-values |
|---|---|---|---|---|---|---|---|---|
| | | | | | Mean | 25%-quantile | 75%-quantile | |
| Drosophila C virus | NC_001834.1 | RNA | 4 | 0 | 2.328 | 1 | 4 | 0.046 |
| Cricket paralysis virus | NC_003924.1 | RNA | 6 | 1 | 3.051 | 1 | 5 | 0.182 |
| Nora virus | NC_007919.3 | RNA | 3 | 1 | 1.418 | 0 | 3 | 0.904 |
| Sigma virus | JX403934.1 | RNA | 4 | 0 | 0.687 | 0 | 2 | 0.932 |
| Acute bee paralysis virus | NC_002548.1 | RNA | 5 | 0 | 2.542 | 1 | 5 | 0.046 |
| Deformed wing virus | NC_004830.2 | RNA | 4 | 0 | 2.187 | 0 | 4 | 0.07 |
| SARS-CoV-2 | NC_045512.2 | RNA | 18 | 1 | 8.817 | 5 | 13 | 0.002 |
| Herpes simplex virus 2 | NC_001798.2 | DNA | 56 | 11 | 11.344 | 6 | 17 | 0.924 |
| Human cytomegalovirus | OK000912.1 | DNA | 116 | 28 | 28.056 | 20 | 37 | 0.918 |
| Kallithea virus | NC_033829.1 | DNA | 43 | 24 | 21.926 | 15 | 28 | 0.422 |
| Invertebrate iridescent virus 6 | NC_003038.1 | DNA | 269 | 158 | 196.661 | 183 | 210 | <0.001 |
| Alphabaculovirus | NC_024625.1 | DNA | 86 | 18 | 30.485 | 22 | 39 | <0.001 |

**Table S6.** Frequencies of deletions in homopolymeric T regions in the Herpes simplex virus serotype 2 genome in sequencing data (accession number: ERR3278849) (53). See separate file.

**Table S7**. List of folding free energy differences in the capsid structure calculated between the wildtype Y95 and its mutants D95, S95, N95, C95 and H95, and for P92T.

| **Variant** | **Avg. energy difference between mutant and wildtype*** | **Total energy of wildtype** | **Avg. energy of mutant** |
|---|---|---|---|
| Tyr95Asp | 0.71 kcal/mol | 1768.31 kcal/mol | 1769.02 kcal/mol |
| Tyr95Ser | 2.1 kcal/mol | 1768.31 kcal/mol | 1770.41 kcal/mol |
| Tyr95Asn | 2.64 kcal/mol | 1768.31 kcal/mol | 1770.95 kcal/mol |
| Tyr95Cys | 3.07 kcal/mol | 1768.31 kcal/mol | 1771.38 kcal/mol |
| Tyr95His | 3.33 kcal/mol | 1768.31 kcal/mol | 1771.64 kcal/mol |
| Pro92Thr | -1.79 kcal/mol | 1768.31 kcal/mol | 1766.52 kcal/mol |
| *ddG= dG(mut) - dG(wildtype) | | | |

### Supplemental Figures

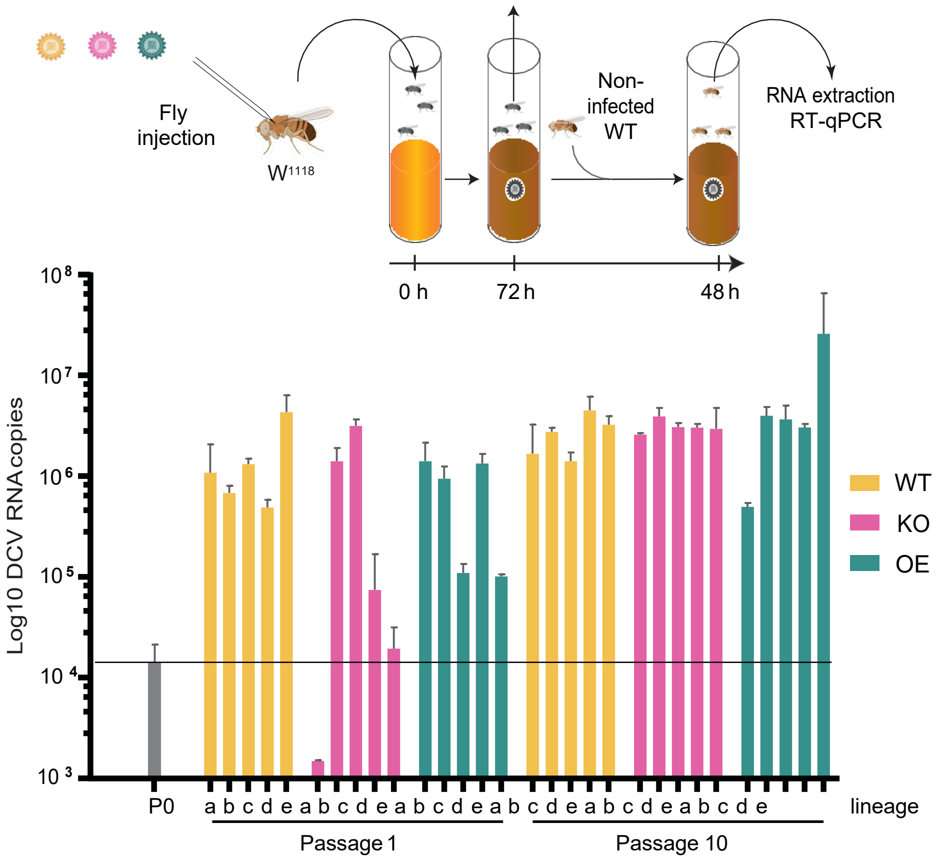

**Figure S1.** Increased replication of evolved virus populations after oral inoculation. 1,000 TCID50 of each virus stock was injected into WT flies and incubated in separate vials for 3 days. Those flies were discarded and uninfected flies were added to the vials and allowed to feed for 2 days. Viral RNA copy numbers were quantified by RT-qPCR from three pools of five flies. The dashed line shows the RNA copies of the parental stock (P0). Results are shown as means and SD of the three replicates of five flies each.

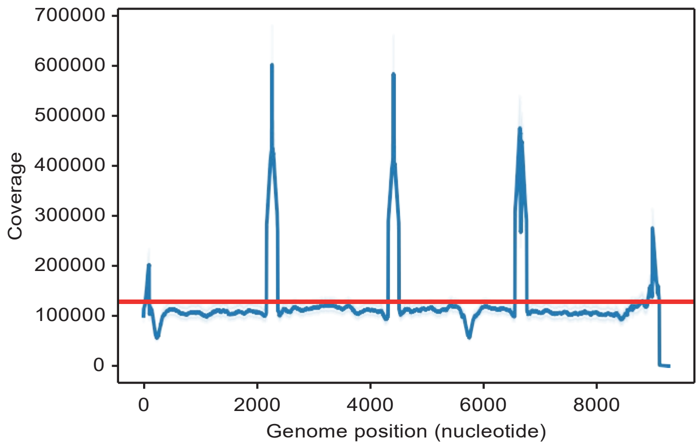

**Figure S2.** Mean sequence coverage across the genome over all experimental lineages and parental stock. The red line indicates the mean coverage (127,542 reads per position), blue shading indicates the 95% confidence intervals based on the bootstrap distribution. Peaks in sequence coverage correspond to regions where amplicons overlap.

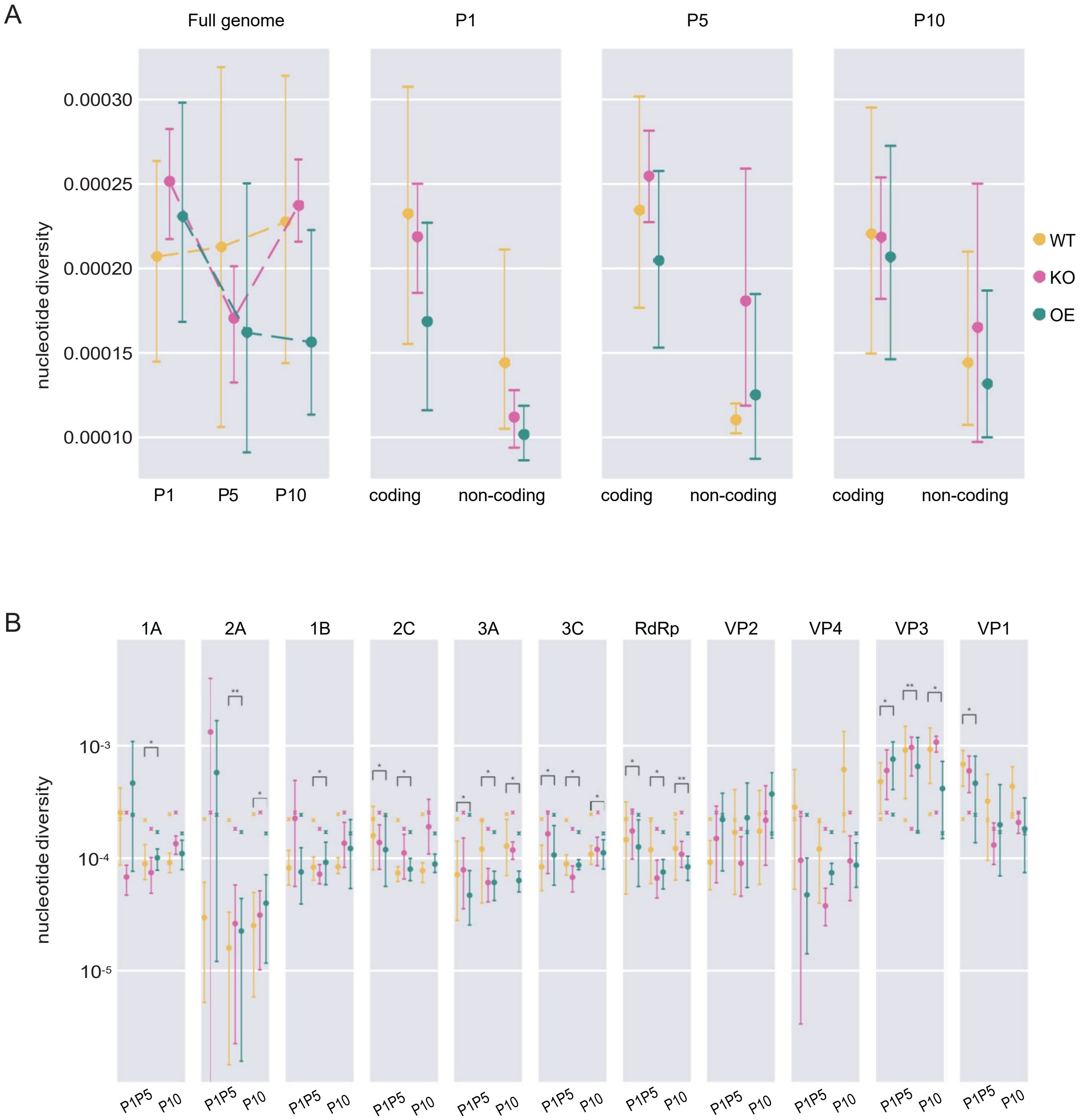

**Figure S3.** Nucleotide diversities of virus populations during serial passage. **(A)** Mean population nucleotide diversity across the full genome (left panel) and in coding and non-coding regions of the genome (right panels) of virus populations evolved in the three host genotypes. Data are shown as means and standard errors across the five lineages for each genotype. **(B)** Mean nucleotide diversity per viral gene marked with dots, the error bars indicating the standard error across the five lineages. Crosses mark the nucleotide diversity across the coding region for each genotype and passage as a baseline. Statistical significance was tested with paired t-tests comparing baseline to the values of each gene. *P*-values are indicated with asterisks: *, $p < 0.05$; **, $p < 0.01$.

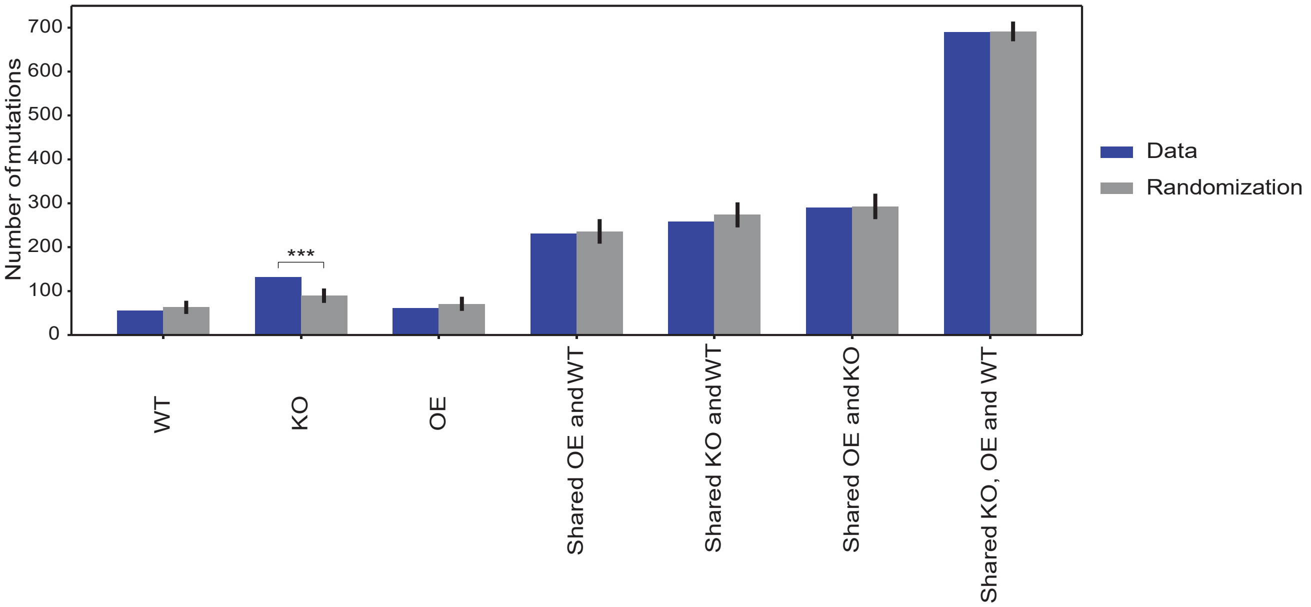

**Figure S4.** Host genotype specific adaptation in *Dcr-2-KO* flies. A permutation test with 1000 randomizations was used to analyze adaptation to specific host genotypes. Label ‘only’ refers to mutations that reach the frequency threshold 0.01% in at least two lineages in the same host genotype. Label ‘shared’ refers to mutations that reach the frequency threshold in more than one genotype. *P*-values are indicated with asterisks: ***, $p < 0.001$. Error bars mark the 95% highest posterior density (HPD) intervals around the mean across the randomizations.

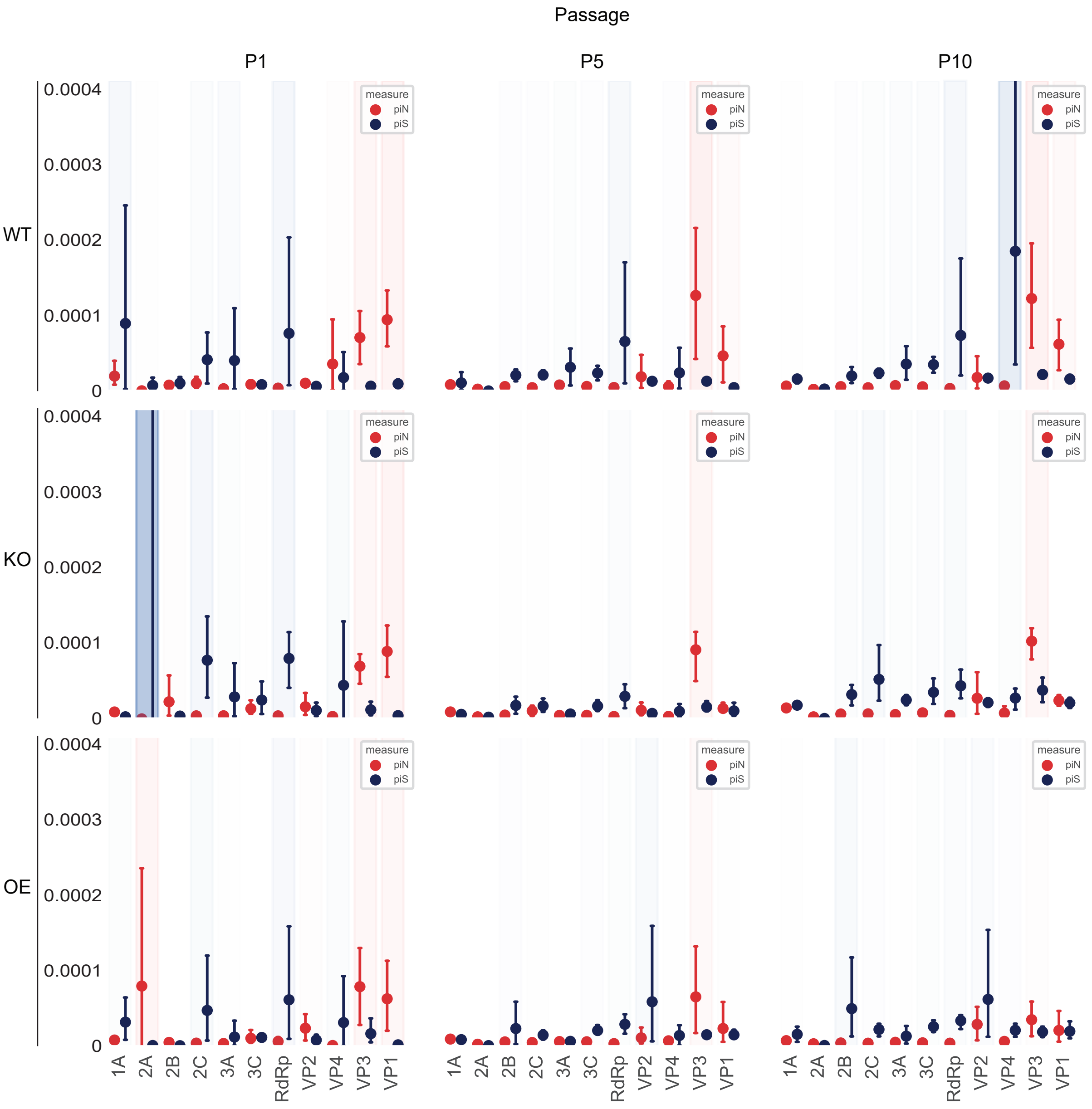

**Figure S5.** Positive selection on the VP3 capsid protein. Comparison of population nucleotide diversity per synonymous (piS) and non-synonymous (piN) site for each viral gene. Red and blue shading indicate piN - piS > 0 and piN - piS < 0, respectively. Error bars indicate 95% confidence intervals based on the bootstrap distribution.

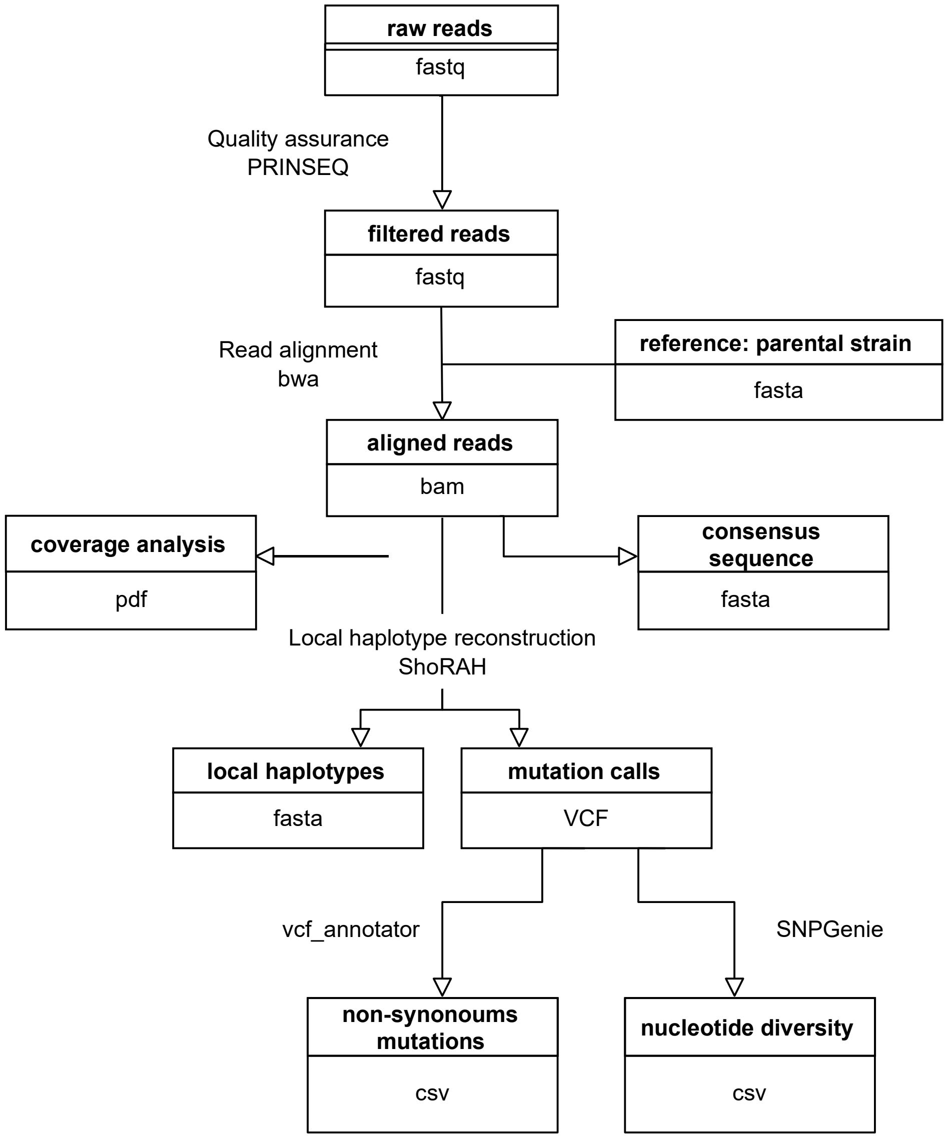

**Figure S6.** Data processing pipeline overview: Quality assurance, Multiple sequence alignment, Coverage analysis, Consensus sequence calling, Mutation and local haplotype calling, Annotation of mutations calls, Computation of nucleotide diversity.

### Supplementary references

1. Longdon B, Day JP, Alves JM, Smith SCL, Houslay TM, McGonigle JE, et al. Host shifts result in parallel genetic changes when viruses evolve in closely related species. PLoS Pathog. 2018 Apr 12;14(4):e1006951.
2. Martinez J, Bruner-Montero G, Arunkumar R, Smith SCL, Day JP, Longdon B, et al. Virus evolution in Wolbachia-infected Drosophila. Proc Biol Sci. 2019 Nov 6;286(1914):20192117.
3. Mongelli V, Lequime S, Kousathanas A, Gausson V, Blanc H, Nigg J, et al. Innate immune pathways act synergistically to constrain RNA virus evolution in Drosophila melanogaster. Nat Ecol Evol. 2022 Mar 10;
4. López-Muñoz AD, Rastrojo A, Kropp KA, Viejo-Borbolla A, Alcamí A. Combination of long- and short-read sequencing fully resolves complex repeats of herpes simplex virus 2 strain MS complete genome. Microb Genom. 2021 Jun;7(6).
